# Far from equilibrium dynamics of tracer particles embedded in a growing multicellular spheroid

**DOI:** 10.1101/2020.03.28.013888

**Authors:** Himadri S. Samanta, Sumit Sinha, D. Thirumalai

## Abstract

Local stresses on the cancer cells (CCs) have been measured by embedding inert tracer particles (TPs) in a growing multicellular spheroid. The utility of the experiments requires that the TPs do not alter the CC microenvironment. We show, using theory and extensive simulations, that proliferation and apoptosis of the CCs, drive the dynamics of the TPs far from equilibrium. On times less than the CC division times, the TPs exhibit sub-diffusive behavior (the mean square displacement, 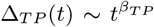 with *β*_*TP*_ < 1). Surprisingly, in the long-time limit, the motion of the TPs is *hyper-diffusive* (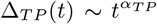 with *α*_*TP*_ > 2) due to persistent directed motion for a number of CC division times. In contrast, CC proliferation randomizes their motion resulting from jamming at short times to super-diffusive behavior, with *α*_*CC*_ exceeding unity, at long times. Surprisingly, the effect of the TPs on CC dynamics and radial pressure is negligible, suggesting that the TPs are reliable reporters of the CC microenvironment.

---

The interplay between short-range forces and non-equilibrium processes arising from cell division and apoptosis gives rise to unexpected dynamics in the collective migration of cancer cells [1–7]. An example is the invasion of cancer cells (CCs) in a growing multicellular spheroid (MCS), which is relevant in cancer metastasis [6, 7]. Imaging experiments show that collective migration of a group of cells that maintain contact for a long period of time exhibits far from equilibrium characteristics [7–11]. Simulations and theory explain some of the experimental observations [12–14]. Dynamics in a growing MCS is reminiscent of the influence of active forces in abiotic systems [15–17]. In a growing MCS the analogue of active forces are self-generated [18], arising from biological events characterized here by cell division and apoptosis.

Experiments that probe the local stresses or pressure on the CCs [19–24] have provided insights into the mechanism by which the CCs invade the extracellular matrix. Recently, the stresses within MCSs were measured [24] by embedding micron-sized inert deformable polyacrylamide beads as probes. For a TP to be a reliable sensor, it should not alter the dynamics of the CCs and the microenvironment, as assessed by local pressure. However, the influence of TPs on the CCs is unknown – a gap that we fill here.

We used theory and simulations to probe the dynamics of the TPs in the MCS, and the impact of the TPs on the CC mobility as well as radial pressure. The central results are : (1) The TPs exhibit sub-diffusive motion in the intermediate time scale 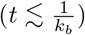, with the mean squared displacement, 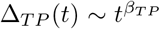 with *β*_*TP*_ less than unity. In the long time limit, 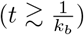, the TPs undergo *hyper-diffusive* motion, 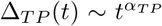 with *α*_*TP*_ greater than 2. In contrast, the CCs, which are jammed at 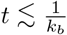, exhibit *super-diffusive* dynamics. (2) The dynamics of the CCs in the intermediate time regime changes slightly as the size of the TPs are varied. Remarkably, the long time exponent (*α*_*CC*_) as well as the decrease in the pressure as a function of the distance from the center of the MCS remains unaffected in the presence of TPs.

## Theory

We consider the dynamics of the TPs in a growing MCS in a dissipative environment by neglecting inertial effects. The deformable but inert TPs, do not grow, divide or undergo apoptosis. The CCs grow and divide at a given rate, and undergo apoptosis (see Figure 1). The CCs and TPs experience systematic short-ranged attractive and repulsive forces arising from other CCs and TPs [12]. To obtain the dynamics of the TPs and CCs, we introduce the density fields *ϕ*(**r**, *t*) = Σ_*i*_*δ*[**r** − **r**_*i*_(*t*)] for the CCs, and *ψ*(**r**, *t*) = Σ_*i*_*δ*[**r** − **r**_*i*_(*t*)] for the TPs. A formally exact Langevin equation for *ϕ*(**r***, t*), and *ψ*(**r***, t*) may be derived using the Dean’s method [25] accounting for diffusion and non-linear interactions.

**Figure 1:**
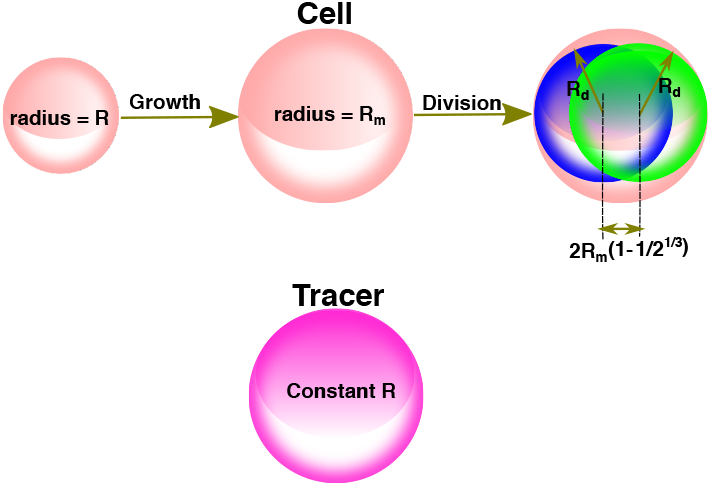
Difference between the CCs (pink sphere) and the TPs (magenta sphere). Cell division creates two daughter CCs (blue and green sphere), each with 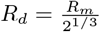 Their relative position is displaced by 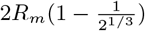 with the orientation being random with respect to the sphere center.

To model a growing MCS, we modify the density equation for the CCs phenomenologically by adding a non-linear source term, ∝ *ϕ*(*ϕ*_0_ − *ϕ*), accounting for cell birth and apoptosis, and a non-equilibrium noise term that breaks the CC number conservation. The noise, *f*_*ϕ*_, satisfies < *f*_*ϕ*_(**r***, t*)*f*_*ϕ*_(**r′**, *t′*) >= *δ*(**r** − **r′**)*δ*(*t* − *t′*). The source term, ∝ *ϕ*(*ϕ*_0_ − *ϕ*), represents the birth and apoptosis, with 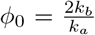 [26,27]. The coefficient, 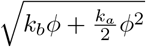 of *f*_*ϕ*_, is the noise strength, corresponding to number fluctuations.

The *ϕ*(**r**, *t*) and *ψ*(**r**, *t*) fields obey,

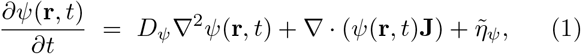

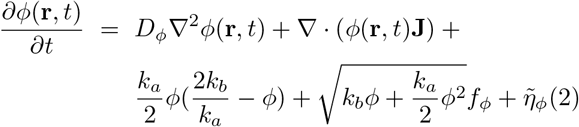

where **J** = *∫*_**r′**_ [*ψ*(**r′**, *t*) + *ϕ*(**r′**, *t*)]∇*U* (**r** − **r′**), 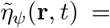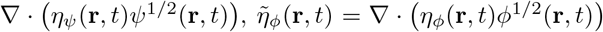, and *η*_*ϕ,ψ*_ satisfies < *η*_*ϕ,ψ*_(**r**, *t*)*η*_*ϕ,ψ*_(**r**′, *t*′) >= *δ*(**r** − **r**′)*δ*(*t* − *t*′). The second term in Eq.(1) accounts for the TP-TP interactions (∇ · *ψ*(**r**, *t*) *∫*_**r′**_ *ψ*(**r′**, *t*)∇*U* (**r** − **r′**))) and TP-CC interactions, (∇·(ψ(**r′**,*t*) *∫*_**r′**_ *ϕ*(**r**, *t*))∇*U*(**r** − **r′**))). The influence of the CCs on the TP dynamics is reflected in the TP-CC coupling. The third term in Eq.(2) results from cell birth and apoptosis, and the fourth term in Eq.(2) is the non-equilibrium noise. Eq. 1 does not satisfy the fluctuation-dissipation theorem in the tumor growth phase.

To obtain the exponents describing the dependence of the mean-square displacement (MSD) on time, we consider a change in scale, **r** → *s***r**, and *t* → *s*^*z*^*t* where *z* is the dynamic exponent. We use the Parisi-Wu [28–30] technique to calculate *z*. The main features of the method, which are not needed to understand the central results here, are relegated to section I in the SI.

## Simulations

We simulated a 3D MCS with embedded TPs using an agent-based model [12, 14, 31]. The cells, treated as interacting soft deformable spheres, grow with time, and divide into two identical daughter cells upon reaching a critical radius (*R*_*m*_). The mean cell cycle time, 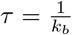 is 15 hrs is used if not mentioned explicitly. The CCs also undergo apoptosis at the rate *k*_*a*_ ≪ *k*_*b*_. The sizes and the number of TPs are held constant.

To determine the TP dynamics, we model the CC-CC, CC-TP, and TP-TP interactions using two potentials (Gaussian and Hertz) to ensure that the qualitative results are robust (simulation details are in section II in the SI).

The equation of motion for the dynamics of TP and CCs is taken to be [12], 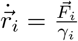 where 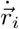 is the velocity of *i^th^* CC or TP, 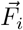 is the force on *i*^*th*^ CC/TP, and *γ*_*i*_ is the damping term. We used a pressure inhibition mechanism to model the observed growth dynamics in solid tumors [12]. Dormancy or the growth phase of the CCs depends on the local microenvironment, determined by the pressure on the *i*^*th*^ cell (Figure 1). Cell division and the placement of the daughter cells add stochasticity in the dynamics of the CCs. We initiated the simulations with 100 TPs and 100 CCs. The spatial coordinates of the CCs and TPs were sampled using a normal distribution with mean zero and standard deviation, 50 *μm*.

### Results

Theory and simulations predict that, in the limit 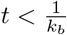, the TPs exhibit sub-diffusive behavior (Table I). The non-linear terms for the TP-TP and TP-CC interactions determine the scaling laws. A power counting analysis shows that the dynamical exponent 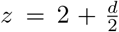. As a consequence, the MSD of a TP behaves as, 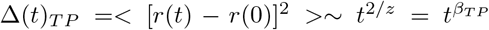. Because *β*_*TP*_ is less than unity we surmise that the TPs are jammed.

**TABLE I:**
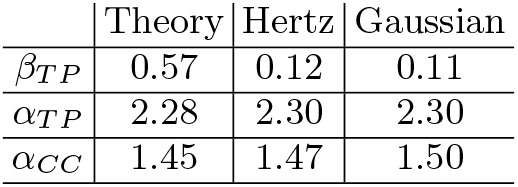
MSD exponents from theory and simulations.

We also calculated Δ_*TP*_(*t*) = 〈[**r**(*t*) − **r**(0)]^2^〉, by averaging over ≈ 2,000 trajectories, for the Gaussian (Figure 2) and Hertz (Figure S2a in the SI) potentials. Similar behavior is found in both cases. The duration of the plateau (Figure 2) increases as the cell cycle time increases. Although jamming behavior predicted theoretically is consistent with the simulations the *β*_*TP*_ exponents differ (Table I).

**Figure 2:**
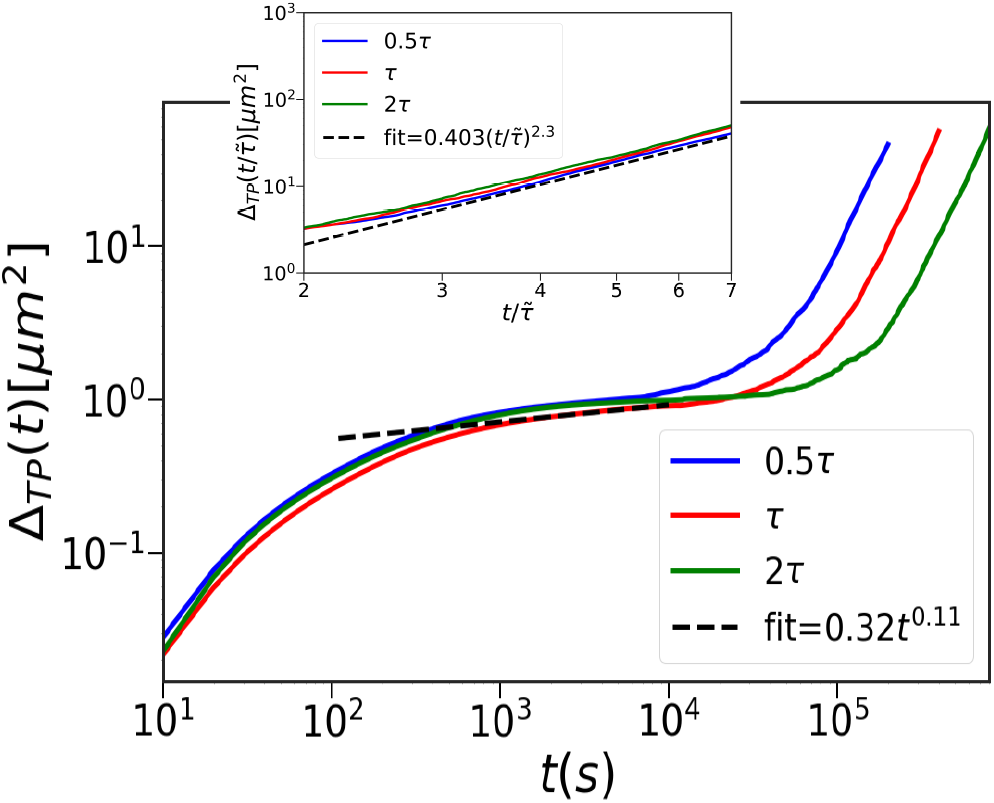
MSD of the TPs (Δ*TP*) using the Gaussian potential. The curves are for 3 cell cycle times (blue (0.5*τ*), red (*τ*), and green (2*τ*). Time to reach the hyper-diffusive behavior, which is preceded by a jamming regime (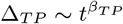, *β*_*TP*_ = 0.11, shown in black dashed line), increases with *τ*. The inset focusses on the hyper-diffusive regime 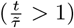. 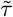 is the cell cycle time for the respective curves. Time is scaled by 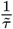. 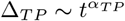, *α*_*TP*_ = 2.3 (dashed black line).

In the 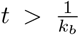 limit, theory and simulations predict hyper-diffusive dynamics, 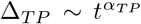 with *α*_*TP*_ > 2. Non-linearities, due to the TP-CC interactions together with cell division and apoptosis, determine the long-time behavior of the TPs. Using the scaling analysis as before, the theory predicts hyper-diffusive dynamics for the TPs (Table I). Variations in *k*_*b*_ do not change the value of *α*_*TP*_. It merely changes the coefficients of the linear term and the value of 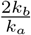. Therefore, *α*_*TP*_ is independent of the cell cycle time in the long time limit. Simulation results are in excellent agreement with the theoretical predictions. For times exceeding the cell cycle time, (*t* > *τ*), the TPs exhibit hyper-diffusive behavior (Figure 2). For the Gaussian potential, we obtain *α*_*TP*_ ≈ 2.3 (see the inset in Figure 2) for 0.5*τ*, *τ* and 2*τ*. Thus, *α*_*TP*_ is independent of cell cycle time. Note that TP-TP interactions play an insignificant role in the dynamics of TPs or the CCs (Figure S3 in the SI). They merely alter the amplitude Δ_*TP*_ in the intermediate time. They do not affect the long time dynamics

Figure 3 shows that changing the tracer size affects only the amplitude of Δ_*CC*_ in the intermediate time without altering the *α*_*TP*_ values (see also figures S4 and S5 in the SI). This is because the TP size only changes the nature of the short-range interactions without introducing any new scale. Since, *α*_*CC*_ is a consequence of the long-range spatial and temporal correlations that emerge because of the inequality *k*_*b*_ ≫ *k*_*a*_, it is independent of the tracer size (section IIIE of SI).

**Figure 3:**
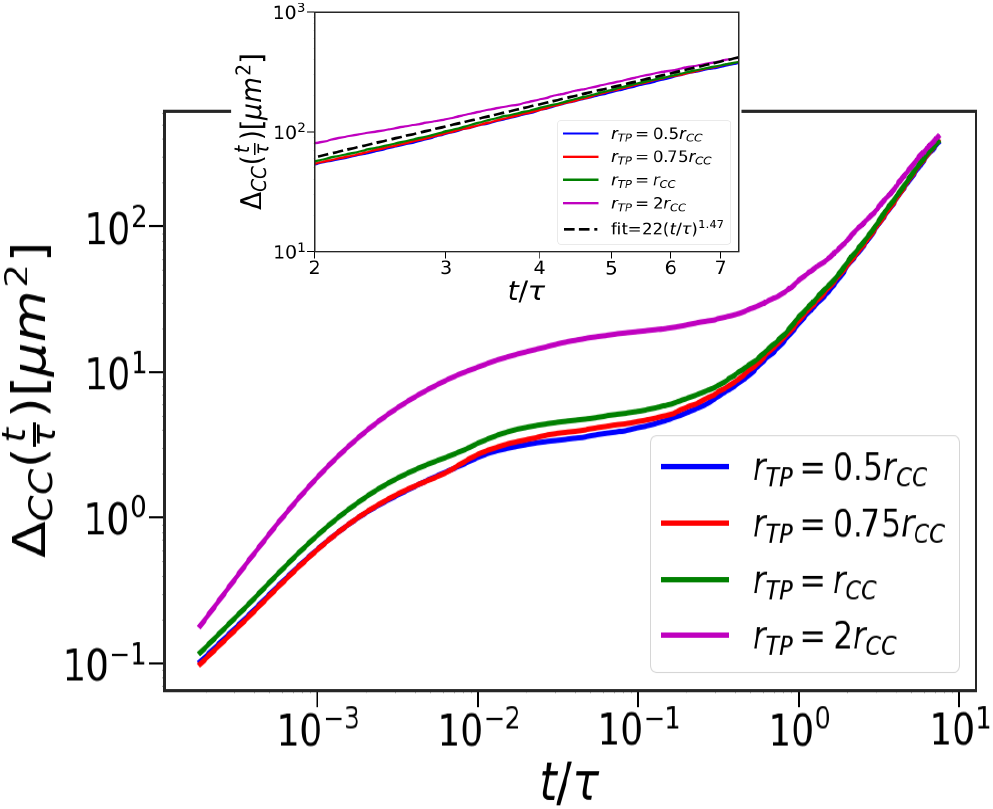
CC dynamics in the presence of the TPs. Δ_*CC*_ using the Hertz potential. The curves are for different TP radius (magenta - *r*_*TP*_ = 2 *r*_*CC*_, green - *r*_*TP*_ = *r*_*CC*_, and red - *r*_*TP*_ = 0.75*r*_*CC*_, and blue - *r*_*TP*_ = 0.5*r*_*CC*_, where *r*_*CC*_ = 4.5*μm* is the average cell radius. Inset shows Δ_*CC*_ for the four curves, focusing on the super-diffusive regime. The black dashed line is drawn with *α*_*CC*_ = 1.47.

In the absence of the TPs, birth and apoptosis determine the CC dynamics in the long time regime, yielding *α*_*CC*_ = 1.33 [12, 13]. When TPs are present, the CCs continues to exhibit super-diffusive motion with a modest increase in *α*_*CC*_ (Table I). The CC pair-correlation, 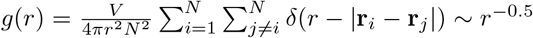, in the presence and absence of TPs at *t* ≈ 8*τ* (Figure 4a). The dynamically-induced CC correlations is independent of the TPs, thus explaining the insignificant effect of the TPs on the CC dynamics.

**Figure 4:**
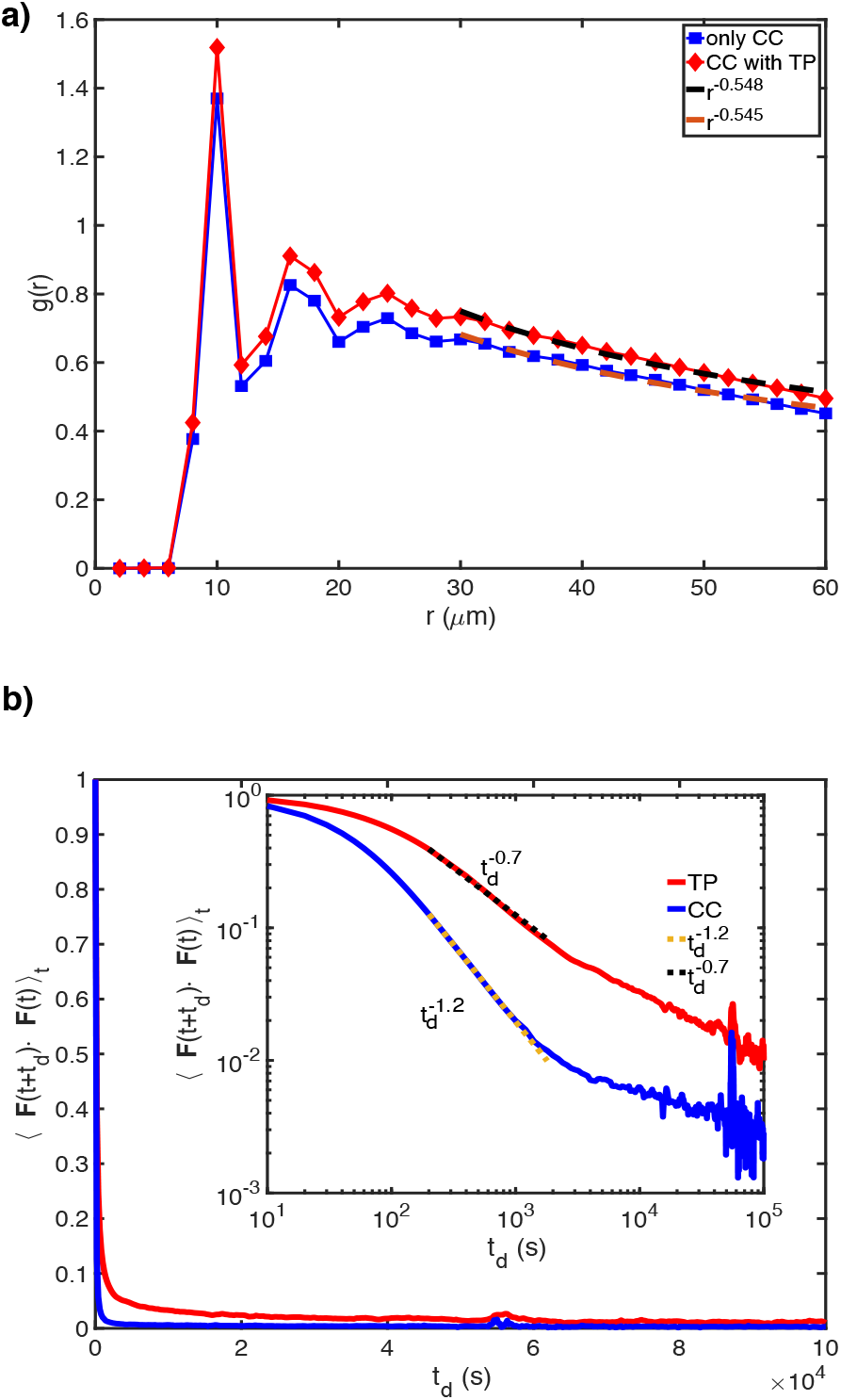
(a) CC correlation function, (*g*(*r*)), as a function of inter-cellular distance. Red (blue) curve shows *g*(*r*) in presence (absence) of TPs. The two dashed lines (black and orange) are power law fits to *g*(*r*) in the large *r* limit. **(b)** Force force autocorrelation (FFA), as a function of delay time (*t*_*d*_). The red (blue) curve shows the FFA for TPs (CCs). Inset shows FFA on log-log scale. The black (yellow) dashed line is a power law fit with exponent of −0.7 (−1.2).

To explain the finding, *α*_*TP*_ > *α*_*CC*_, we calculated the force-force autocorrelation function, FFA= 〈**F**(*t*+*t*_*d*_) · **F**(*t*)〉_*t*_. Here, 〈…〉_*t*_ is the time average and *t*_*d*_ is a delay time. Since, the TPs (CCs) exhibit hyper-diffusion (super-diffusion), we expect that the FFA of TPs should decay slower relative to the CCs, which is confirmed in Figure 4b, which shows that the FFA for the TPs (CCs) decays as 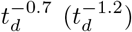. Thus, the TP motion is significantly more persistent than the CCs, which explains the hyper-diffusive nature of the TPs.

A mechanistic explanation for *α*_*TP*_ > *α*_*CC*_ can be gleaned from the simulations. To illustrate the difference between the dynamics of the TPs and CCs, we calculated the angle (*θ*) between two consecutive time steps in a trajectory. We define 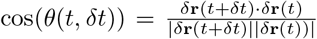, where *δ***r**(*t*) = **r**(*t*+*δt*)−**r**(*t*). Figure 5a shows the ensemble and time averaged cos *θ* distribution for 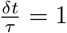. If the motion of the CCs and TPs were diffusive, the distribution of cos *θ* would be uniformly distributed from −1 to 1. However, Figure 5a shows that *P* (cos *θ*) is skewed towards unity implying that the motion of both the TPs and CCs are persistent. Interestingly, the skewness is more pronounced for the TPs compared to the CCs (Figure 5a). During every cell division, the motion of CCs is randomized and hence the persistence is small compared to TPs. This explains the hyper-diffusive (super-diffusive) for the TPs (CCs) and thus *α*_*TP*_ > *α*_*CC*_ in the long time limit (see Figure S6 which shows TPs move much more straighter than CCs).

**Figure 5:**
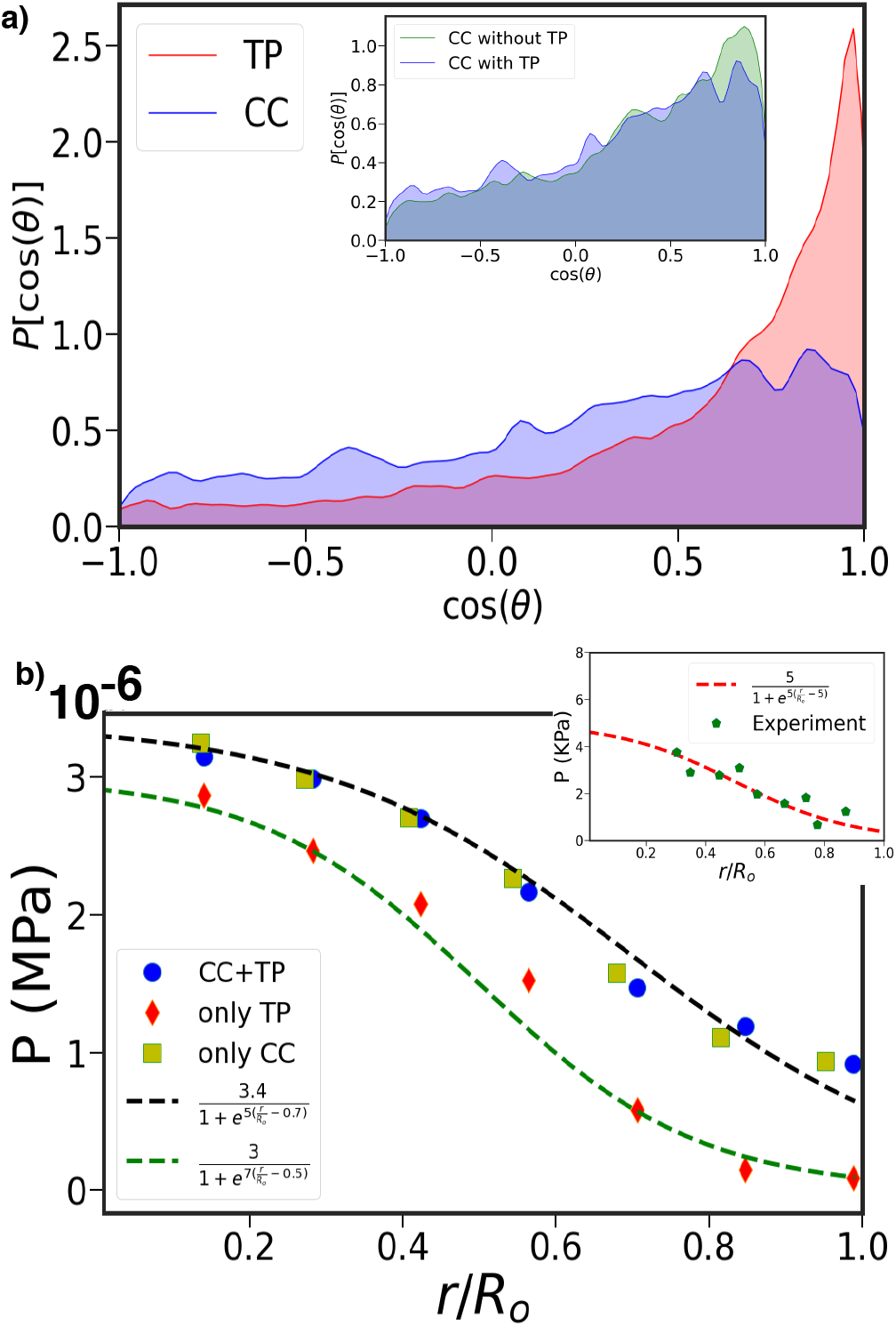
(a) Distribution of cos(*θ*) for the TPs and CCs. *θ* is the angle between two consecutive steps along a trajectory. The red (blue) plot is for the TPs (CCs). The distribution is skewed to cos(*θ*) > 0, indicative of persistent motion. Extent of skewness is greater for TPs than CCs. The inset shows the *P* [cos(*θ*)] of CCs with and without TPs. **(b)** Pressure as a function of the radial distance (*r* scaled by *R*_0_: tumor radius ≈ 100*μm*) from tumor center. The blue circles correspond to local pressure measured using both CCs and TPs. The red data points is local pressure measured using just the TPs using simulations containing both the CCs and TPs. The greenish yellow squares give the local pressure obtained in simulations with only the CCs. The black and green dashed lines is logistic fit. The inset shows the radial pressure profile in experiments (green data points). The red dashed lines are logistic fits. Note that the magnitude of pressure measured in simulations and experiments are in qualitative agreement.

To assess the effectiveness of the TPs as sensors of the local microenvironment, we calculated the radial pressure (Eq S28 in the SI) profile at *t* = 7.5*τ*, which has been measured [24]. We measured pressure profiles using Eq. S28 (in the SI) for the system with both CCs and TPs. We also calculated pressure experienced just by the TPs. Finally, we computed pressure in a system consisting of solely the CCs. Two pertinent remarks about Figure (5b are worth making. (i) All three curves qualitatively capture the pressure profile found in the experiments. Pressure decreases roughly by factor of four, as the distance *r* from the center of the tumor increases. The pressure is almost constant in the core, with a decrease that can be fit using the logistic function, as the boundary of the tumor is reached (Figure (5b). The high core pressure is due to small number of cell divisions. As a consequence, the CCs are jammed, leading to high internal pressure. As *r* increases, the CCs proliferate, resulting in a decrease in self generated stress, and consequently a decreases in the pressure (Figure (5b).(ii) Most importantly, the CC pressure profiles are unaltered even in the presence of the TPs (compare blue circles and greenish yellow squares in Figure (5b), which shows that the latter does faithfully report the microenvironment of the CCs.

We conclude with a few comments. (i) The excellent agreement for *α*_*TP*_ between simulations and theory suggests that the CC dynamics is determined by the overall tumor growth, and not the details of the TP-CC interactions. Although *α*_*TP*_ > *α*_*CC*_, the CCs move a larger distance than the TPs (compare Figure 2 and Figure 3), which implies that the effective diffusion constant of the CCs is greater than the TPs. (ii) The radial dependence of the pressure (Figure 5b), with the core experiencing higher values than the periphery cells, is another manifestation of non-equilibrium that is induced by an imbalance in division and apoptosis rates of the CCs. (iii) Most importantly, we find that the long time dynamics of CCs and the pressure profiles are unchanged when the TPs are embedded in the MCS. Therefore, it is justified to use the TPs as tumor stress sensors.

## Acknowledgements

We thank Mauro Mugnai, Hung Nguyen, Rytota Takaki and Davin Jeong for useful discussions and comments on the manuscript. This work is supported by the National Science Foundation (PHY 17-08128), and the Collie-Welch Chair through the Welch Foundation (F-0019).

