## Supplementary Information for "Far from equilibrium dynamics of tracer particles embedded in a growing multicellular spheroid"

(Dated: October 14, 2020)

#### Contents


I. Theory 2
  A. Time-dependent equations for TP and CC densities 2
  B. Stochastic Quantization 5
  C. Theory for TP and CC dynamics 7

II. Simulation Models: 8
  A. Hertz potential 9
  B. Gaussian potential 9
  C. Equation of Motion 10
  D. Cell division and Apoptosis 10

III. Details of the Results 11
  A. TPs exhibit sub-diffusive dynamics in the intermediate time regime 11
  B. Hyper-diffusion of the TPs in the long time limit 11
  C. k-dependent diffusion coefficients 11
  D. $\alpha_{TP}$ is nearly independent of the TP size 13
  E. Influence of the TPs on CC dynamics 13
  F. Straigtness Index 17

References 18


### I. THEORY

#### A. Time-dependent equations for TP and CC densities

Let us consider the dynamics of the tracer particles (TPs) in a growing tumor spheroid. The TPs experience systematic short-range interactions due to volume excluded from the neighboring TPs and the cancer cells (CCs). In addition, they are also subject to a random force characterized by a Gaussian white noise spectrum. For mathematical convenience, the inter-cell interactions are modeled as a sum of attractive, and repulsive excluded volume interactions. We assume that the dynamics of the system, consisting of the CCs and TPs (see Figure S1 for snapshots generated in simulations) can be described by the overdamped Langevin equation,

$$\frac{d\mathbf{r}_i}{dt} = -\sum_{j=1}^{N} \nabla U(|\mathbf{r}_i - \mathbf{r}_j|) + \boldsymbol{\eta}_i(t), \tag{S1}$$

where $\mathbf{r}_i$ is the position of a CC or a TP, and $\boldsymbol{\eta}_i(t)$ is a Gaussian random force with white noise spectrum. To keep the problem theoretically tractable, the form of $U(|\mathbf{r}_i - \mathbf{r}_j|)$ between a pair of particles (can be either TP-TP, TP-CC or CC-CC) is taken to be ,

$$U(|\mathbf{r}(i) - \mathbf{r}(j)|) = \frac{\nu}{(2\pi\lambda^2)^{3/2}} e^{\frac{-|\mathbf{r}(i)-\mathbf{r}(j)|^2}{2\lambda^2}} - \frac{\kappa}{(2\pi\sigma^2)^{3/2}} e^{\frac{-|\mathbf{r}(i)-\mathbf{r}(j)|^2}{2\sigma^2}}, \tag{S2}$$

where, $\lambda$ and $\sigma$ are the ranges of the repulsive and attractive interactions, and $\nu$ and $\kappa$ are the interaction strengths. Thus, the interactions involving the mixture of CCs and TPs are identical.

When equation S1 is used to characterize the dynamics of the TPs, the potential $U_{TP}$ contains both the TP-TP and TP-CC interactions with the corresponding attractive (repulsive) interaction ranges being $\sigma_1(\lambda_1)$ and $\sigma_2(\lambda_2)$, respectively. The potential $U_{CC}$ for the CCs mimics cell-cell adhesion (second term in the above equation) and excluded volume interactions, and the CC-TP interactions.

A closed form Langevin equation for the CC density, $\phi(\mathbf{r}, t) = \sum_i \phi_i(\mathbf{r}, t)$, where $\phi_i(\mathbf{r}, t) = \delta[\mathbf{r} - \mathbf{r_i(t)}]$, can be obtained using the method introduced elsewhere [1]. The time evolution of $\phi(\mathbf{r}, t)$ is given by,

$$\begin{aligned}\frac{\partial \phi(\mathbf{r}, t)}{\partial t} &= \nabla \cdot \left( \phi(\mathbf{r}, t) \int_{\mathbf{r}'} [\psi(\mathbf{r}', t) \nabla U_{CC-TP}(\mathbf{r} - \mathbf{r}') + \phi(\mathbf{r}', t) \nabla U_{CC}(\mathbf{r} - \mathbf{r}')] \right) \\ &+ D_\phi \nabla^2 \phi(\mathbf{r}, t) + \nabla \cdot \left( \eta_\phi(\mathbf{r}, t) \phi^{1/2}(\mathbf{r}, t) \right),\end{aligned} \tag{S3}$$

where $\eta_\phi$ satisfies $< \eta_\phi(\mathbf{r}, t)\eta_\phi(\mathbf{r}', t') >= 2D_\phi \delta(\mathbf{r} - \mathbf{r}')\delta(t - t')$. Similarly, the evolution of the density function for a single TP, $\psi(\mathbf{r}, t) = \sum_i \psi_i(\mathbf{r}, t) = \sum_i \delta[\mathbf{r} - \mathbf{r_i(t)}]$, may be written as,

$$\begin{aligned}\frac{\partial \psi(\mathbf{r}, t)}{\partial t} &= D_\psi \nabla^2 \psi(\mathbf{r}, t) + \nabla \cdot \left( \psi(\mathbf{r}, t) \int_{\mathbf{r}'} [\psi(\mathbf{r}', t) \nabla U_{TP}(\mathbf{r} - \mathbf{r}') \right. \\ &\left. + \phi(\mathbf{r}', t) \nabla U_{TP-CC}(\mathbf{r} - \mathbf{r}')] \right) + \nabla \cdot \left( \eta_\psi(\mathbf{r}, t) \psi^{1/2}(\mathbf{r}, t) \right).\end{aligned} \tag{S4}$$

where $\eta_\psi$ satisfies $< \eta_\psi(\mathbf{r}, t)\eta_\psi(\mathbf{r}', t') >= 2D_\psi \delta(\mathbf{r} - \mathbf{r}')\delta(t - t')$.

We modify the density evolution for the CCs phenomenologically by adding a source term describing cell division and apoptosis, and a noise term that breaks the CC number conservation. These terms can be formally derived as follows. Birth and apoptosis reactions are given by $X \xrightarrow{k_b} X + X$ and $X + X \xrightarrow{k_a/\Omega} X$, where $k_a/\Omega$ is the apoptosis rate of distinct pairs of cells, and $\Omega$, the volume, will eventually be set to infinity. Let the $\rho$ be the density of cells. The master equation for the evolution of the probability $P_n(t)$ is given by,

$$\frac{dP_n(t)}{dt} = k_b(n-1)P_{n-1} - k_b n P_n - \frac{k_a}{\Omega}\frac{n(n-1)}{2}P_n + \frac{k_a}{\Omega}\frac{n(n+1)}{2}P_{n+1} \tag{S5}$$

Eq.(S5) can be expanded up to second order in $n$ and the resulting equation is given by

$$\begin{aligned}\frac{dP_n(t)}{dt} &= k_b(1 - \frac{\partial}{\partial n} + \frac{1}{2}\frac{\partial^2}{\partial n^2})(n)P_n - k_b n P_n \\ &- \frac{k_a}{\Omega}\frac{n(n-1)}{2}P_n + \frac{k_a}{\Omega}(1 - \frac{\partial}{\partial n} + \frac{1}{2}\frac{\partial^2}{\partial n^2})\frac{n(n-1)}{2}P_n \\ &= \frac{\partial}{\partial n}[-k_b n + \frac{k_a}{2\Omega}n(n-1)]P_n(t) + \frac{\partial^2}{\partial n^2}[\frac{k_b}{2}n + \frac{k_a}{4\Omega}n(n-1)]P_n(t)\end{aligned} \tag{S6}$$

The corresponding Langevin equation for the density $\rho = n/\Omega$ is,

$$\frac{\partial\rho(t)}{\partial t} = k_b\rho(t) - \frac{k_a}{2}\rho(t)(\rho(t) - \frac{1}{\Omega}) + \frac{1}{\sqrt{2\Omega}}\sqrt{k_b\rho(t) + \frac{k_a}{2}\rho(t)(\rho(t) - \frac{1}{\Omega})}f(t) \tag{S7}$$

In order to account for spatial variations, we generalize this scheme by considering the concentration in the $i$th volume element,

$$\frac{\partial\rho_i(t)}{\partial t} = k_b\rho_i(t) - \frac{k_a}{2}\rho_i(t)(\rho_i(t) - \frac{1}{\Omega}) + \frac{1}{\sqrt{2\Omega}}\sqrt{k_b\rho_i(t) + \frac{k_a}{2}\rho_i(t)(\rho_i(t) - \frac{1}{\Omega})}f_i(t), \tag{S8}$$

where $< f_i(t)f_j(t') >= 2\delta_{ij}\delta(t-t')$. In the continuum limit, $\rho_i \to \rho(\mathbf{r},t)$, $f_i(t) \to f(\mathbf{r},t)$, and $\delta_{ij} \to \Omega\delta(\mathbf{r}-\mathbf{r}')$, we get the following equation,

$$\begin{aligned}\frac{\partial\rho(\mathbf{r},t)}{\partial t} &= (k_b + \frac{k_a}{2\Omega})\rho(\mathbf{r},t) - \frac{k_a}{2}\rho(\mathbf{r},t)^2 + \sqrt{(k_b - \frac{k_a}{2\Omega})\rho(\mathbf{r},t) + \frac{k_a}{2}\rho(\mathbf{r},t)^2}f(\mathbf{r},t) \\ &= \frac{k_a}{2}\rho(\mathbf{r},t)((\frac{2k_b}{k_a} + \frac{1}{\Omega}) - \rho(\mathbf{r},t)) + \sqrt{(k_b - \frac{k_a}{2\Omega})\rho(\mathbf{r},t) + \frac{k_a}{2}\rho(\mathbf{r},t)^2}f(\mathbf{r},t),\end{aligned} \tag{S9}$$

$< f(\mathbf{r},t)f(\mathbf{r}',t') >= \delta(\mathbf{r}-\mathbf{r}')\delta(t-t')$.

By letting $\Omega \to \infty$, the time evolution of the density $\rho$ becomes,

$$\frac{\partial\rho(\mathbf{r},t)}{\partial t} = \frac{k_a}{2}\rho(\mathbf{r},t)\left(\frac{2k_b}{k_a} - \rho(\mathbf{r},t)\right) + \sqrt{k_b\rho(\mathbf{r},t) + \frac{k_a}{2}\rho(\mathbf{r},t)^2}f(\mathbf{r},t). \tag{S10}$$

We add the terms on the right hand side in the density equation for the CCs.

The final Langevin equation, for the time-dependent changes in $\phi(\mathbf{r},t)$ is [2],

$$\begin{aligned}\frac{\partial\phi(\mathbf{r},t)}{\partial t} &= D_\phi\nabla^2\phi(\mathbf{r},t) + \nabla\cdot\Big(\phi(\mathbf{r},t)\int d\mathbf{r}'[\psi(\mathbf{r}',t)\nabla U_{CC-TP}(\mathbf{r}-\mathbf{r}') \\ &+\phi(\mathbf{r}',t)\nabla U_{CC}(\mathbf{r}-\mathbf{r}')]\Big) + \frac{k_a}{2}\phi(\frac{2k_b}{k_a} - \phi) + \nabla\cdot\left(\eta_\phi(\mathbf{r},t)\phi^{1/2}(\mathbf{r},t)\right) + \sqrt{k_b\phi + \frac{k_a}{2}\phi^2}f_\phi\,,\end{aligned} \tag{S11}$$

where $f_\phi$ satisfies $< f_\phi(\mathbf{r},t)f_\phi(\mathbf{r}',t') >= \delta(\mathbf{r}-\mathbf{r}')\delta(t-t')$. The source term $\propto \phi(\phi_0 - \phi)$, represents the cell division and apoptosis, with $\phi_0 = \frac{2k_b}{k_a}$ [3, 4]. The coefficient of $f_\phi$, given by $\sqrt{k_b\phi + \frac{k_a}{2}\phi^2}$, is the strength of the noise corresponding to number fluctuations of the CCs, and is a function of the CC density.

To simplify the multiplicative noise terms (last term in Eq. (S4) and the last two terms in Eq. (S11)), we assume that the density fluctuates around a constant value. We write $\Psi(\mathbf{r},t) = \Psi_0 + \Psi_1(\mathbf{r},t)$ ($\Psi(\mathbf{r},t)$ can be either $\phi$ or $\psi$), and expand Eq. (S4) and Eq. (S11) in terms of $\Psi_1$ up to the lowest order in nonlinearity. We consider the scaling behavior under a change of scale given by $\mathbf{r} \to s\mathbf{r}$ and $t \to s^z t$ with $z$ being the dynamic exponent.

The interplay between adhesion interactions and stochastic birth-apoptosis processes gives rise to long-range correlations in the density fluctuations of the TPs and CCs. We consider the scaling of the correlation of density fluctuations for TP as $< \psi_1(\mathbf{r}', t)\psi_1(\mathbf{r}, t) >\sim |\mathbf{r'} - \mathbf{r}|^{\mathbf{2}\chi_1}$, and similarly for CC, $< \phi_1(\mathbf{r}', \mathbf{t})\phi_1(\mathbf{r}, t) >\sim |\mathbf{r'} - \mathbf{r}|^{\mathbf{2}\chi_2}$, where $\chi_1$ and $\chi_2$ are the exponents corresponding to TP and CC density fluctuations respectively. For $t \lesssim {k_b}^{-1}$, the scaling of the dynamical observables for the TPs is governed by the TP-TP and TP-CC interactions. In Fourier space, the non-linear term $(q.(k-q)\psi_1(q)\psi_1(k-q))$ for the TP-TP interactions scales (Eq. (S4)) as $q^{2-2\chi_1}$. Similarly, the non-linear term $(q.(k-q)\phi_1(q)\psi_1(k-q))$ for the TP-CC interactions scales as $q^{2-\chi_1-\chi_2}$. The degree of non-linearity for both the TP-TP and TP-CC interactions is $q^{2-2\chi_1}$, by noting that $\chi_1 \sim \chi_2$. Therefore, both TP-CC and TP-TP interactions exhibit the same scaling for the dynamical properties of TPs in the $t \lesssim {k_b}^{-1}$ limit. At long times ($t > \frac{1}{k_b}$), the birth-apoptosis term $\phi_1(q)\phi_1(k-q)$ (scales as $q^{-2\chi_2}$), dominates over the short-range CC-CC interactions $q.(k-q)\phi_1(q)\phi_1(k-q)$ ( scales as $q^{2-2\chi_2}$). Therefore, when $t > \frac{1}{k_b}$, the birth-apoptosis non-linearity determines the scaling properties for the TPs through the TP-CC interactions.

##### B. Stochastic Quantization

A classical Langevin equation may be written as a path integral with an appropriate action. Similarly, the path integral given by the distribution,

$$\exp[-\mathcal{S}(\Phi(x,t)]/\int D\Phi \exp[-\mathcal{S}(\Phi(x,t)], \tag{S12}$$

can be described as the equilibrium limit of a statistical system that is coupled to a thermal reservoir [5–8]. The coupling to the thermal bath is simulated by letting $\Phi(x,t)$ ($t$ is real time) evolve in a fictitious time $\tau$ in the presence of stochastic noise subject to a systematic force $\propto \frac{\partial\Phi(x,t,\tau)}{\partial\tau}$. As $\tau \to \infty$, the distribution in the above equation would be recovered provided the stochastic noise satisfies the Fluctuation Dissipation Theorem (FDT). Here, $\mathcal{S}$ is a functional of the collective time-dependent density fields associated with the TPs and CCs.

Consider the evolution of $\Phi(x,t,\tau)$ given by the Langevin equation,

$$\frac{\partial\Phi(x,t,\tau)}{\partial\tau} = -\frac{\delta\mathcal{S}}{\delta\Phi} + g(x,t,\tau) \tag{S13}$$

where $g$ is a Gaussian random variable with zero mean and,

$$< g(x,t,\tau)g(x',t',\tau') >= 2\delta(x-x')\delta(t-t')\delta(\tau-\tau'). \tag{S14}$$

Properties of interest involving the density field, $\Phi(x,t,\tau)$, can be evaluated by averaging over the Gaussian noise $g$, and taking the limit of $\tau \to \infty$. For example,$<$

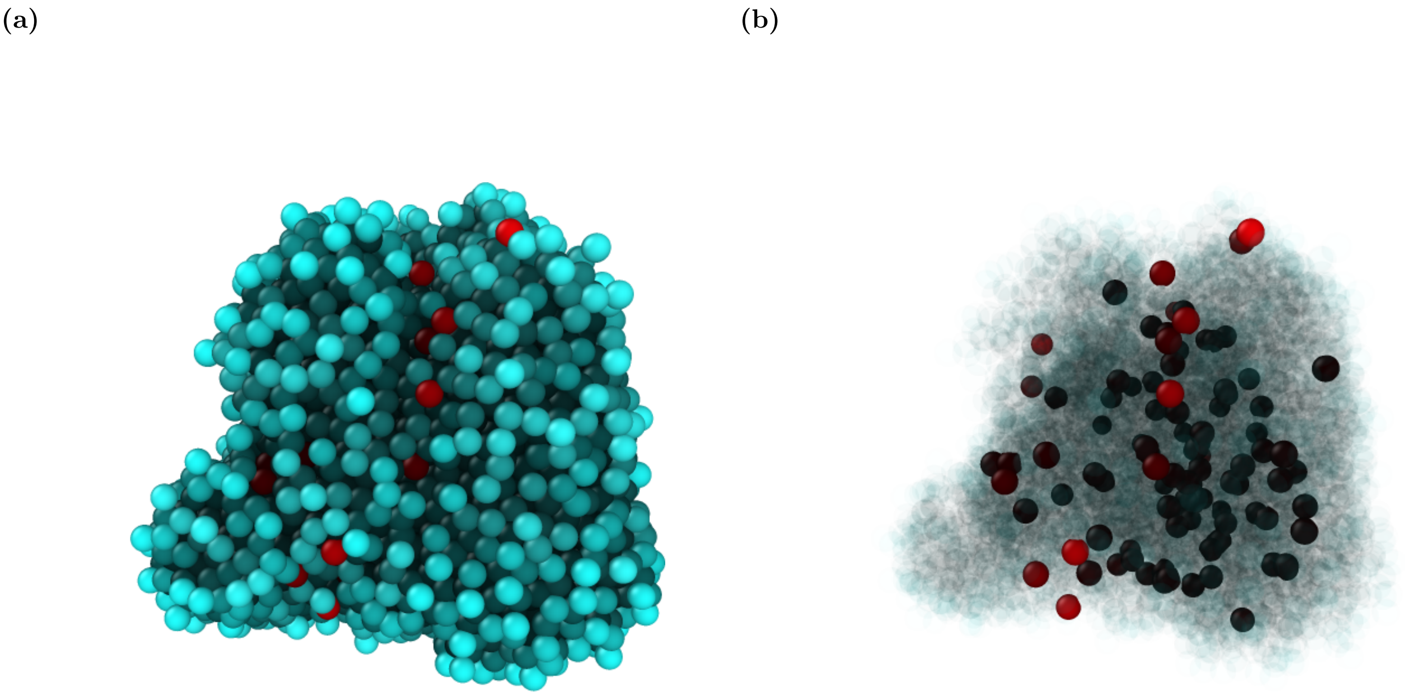

FIG. S1: Snapshots of tumor simulation with embedded tracers. **(a)** A 3D simulated spheroid consisting of approximately 4,800 CCs and 100 TPs. The CCs are in cyan, and the TPs are in red. **(b)** The spheroid shown above was rendered by making the CC cells transparent (light colored cyan) in order to show the interior of the spheroid. The TPs are opaque. Some of the TPs appear as black because it is a depiction of a 3D image. The purpose of displaying these snapshots is to show pictorially that the TPs are randomly distributed within the multicellular spheroid, implying that their migration is largely determined by the forces arising from the CCs.

$\Phi(x_1,t_1)\Phi(x_2,t_2) >$ can obtained using,

$$\lim_{\tau\to\infty} < \Phi(x_1,t_1,\tau)\Phi(x_2,t_2,\tau) >_g \; = \; < \Phi(x_1,t_1)\Phi(x_2,t_2) > \; = \frac{\int \mathcal{D}[\Phi]\Phi(x_1,t_1)\Phi(x_2,t_2))e^{-S[\Phi]}}{\int \mathcal{D}[\Phi]e^{-S[\Phi]}} \tag{S15}$$

We outline the steps to obtain the dynamic exponent $z$.

1. We consider a scalar field $\Phi(k)$ in the momentum space satisfying the general equation of motion,

$$\dot{\Phi}(k) = -A(k)\Phi(k) - B(\Phi) + \eta_\Phi(k), \tag{S16}$$

where $B(\Phi)$ is a non-linear function of $\Phi$ and $\eta_\Phi(k)$ is a noise term with correlation $<\eta_\Phi(k)\eta_\Phi(k')>= 2D_0\delta(k+k')$. The probability distribution corresponding to the noise is given by, $P(\eta_\Phi) \propto \exp[-\int \frac{d^Dk}{(2\pi)^D}\frac{dw}{2\pi}\frac{1}{4D_0}\eta_\Phi(k,w)\eta_\Phi(-k,-w)]$. In the momentum-frequency space, Eq.(S16) reads

$$[-iw + A(k)]\Phi(k,w) + B_{k,w}(\Phi) = \eta_\Phi(k,w) \tag{S17}$$

2. The probability distribution written in terms of $\Phi(k,w)$ instead of $\eta_\Phi(k,w)$ is,

$$\begin{aligned} P &\propto \exp\{-\frac{1}{4D_0}\int_{k,w}\{[-iw + A(k)]\Phi(k,w) + B_{k,w}(\Phi)\}\{[iw + A(-k)]\Phi(-k,-w) + B_{-k,-w}(\Phi)\}\} \\ &= \int \mathcal{D}[\Phi]e^{-S(\Phi)} \end{aligned} \tag{S18}$$

3. Introduce a fictitious time $'\tau'$ and consider all the variables to be functions of $\tau$ in addition to $k$ and $w$. A Langevin equation in the $'\tau'$ variable is,

$$\frac{\partial\Phi(k,w,\tau)}{\partial\tau} = -\frac{\delta S}{\delta\Phi(-k,-w,\tau)} + g(k,w,\tau) \tag{S19}$$

with $< gg >= 2\delta(k+k')\delta(w+w')\delta(\tau-\tau')$.

4. In order to obtain the scaling laws for the mean square displacement (MSD) it suffices to work at arbitrary $\tau$. From Eq.(S19), the Greens function $G$, is

$$G^{-1} = -i\omega_\tau + \frac{w^2 + A^2}{2D_0} + \Sigma(k,w,\omega_\tau), \tag{S20}$$

where $\omega_\tau$ is the frequency corresponding to the fictitious time $\tau$. The nonlinear terms in $B$ contribute to the self-energy $\Sigma(k,w,\omega_\tau)$ . The correlation function is given by $C = \frac{1}{\omega_\tau}ImG$ through the FDT.

5. The nonlinear terms are treated using perturbation theory. A self consistent calculation yields the scaling exponents. The dimensionality of $\Sigma$ can be counted by using the dimensionality of $C$ through FDT relation $C = 1/\omega_\tau ImG$. If the frequency scale is modified to $k^z$, then $G$ scales as $k^{-2z}$. We obtain the dynamic exponent $z$ by matching the power count of last two terms in Eq. (S20).

##### C. Theory for TP and CC dynamics

The evolution of the CC density, described by Eq. (S11) is an out of equilibrium process characterized by the absence of FDT. The usual analytic route to solve the problem is to introduce a response field $\tilde{\phi}$. Here, we need to calculate both the response function ($G =< \phi\tilde{\phi} >$) and correlation function ($C =< \phi\phi >$) separately because of the FDT is violated.

The density and noise fields are functions of $\tau$ in addition to the real parameters, $\mathbf{r}$ and $t$. The equation of motion for $\Psi(\mathbf{r}, t)$ in the '$\tau$' variable is,

$$\frac{\partial \Psi(\mathbf{k}, w, \tau)}{\partial \tau} = -\frac{\delta \mathcal{S}}{\delta \Psi(-\mathbf{k}, -w, \tau)} + g(\mathbf{k}, w, \tau)\,, \tag{S21}$$

with $< gg >= 2\delta(k+k')\delta(w+w')\delta(\tau-\tau')$. The field $\Psi$ represents both $\psi_1$ and $\phi_1$ fields. We are assured that the distribution for the density fields in Eq. S21 will approach $\exp[-\mathcal{S}(\phi_1, \psi_1)]$ in the $\tau \to \infty$ limit, because FDT is preserved in the $\tau$ variable. The action $\mathcal{S}(\phi_1, \psi_1)$ may be obtained by writing the joint probability distribution $P(\eta'_{\phi_1}, \eta'_{\psi_1}) \propto \exp[-\int \frac{d^d\mathbf{k}}{(2\pi)^d}\frac{dw}{2\pi}\{\frac{1}{2(k_a\phi_0+k_b\phi_0^2+D_\phi\phi_0 k^2)}\eta'_{\phi_1}(\mathbf{k}, w)\eta'_{\phi_1}(-\mathbf{k}, -w) + \frac{1}{(2D_\psi\psi_0 k^2)}\eta'_{\psi_1}(\mathbf{k}, w)\eta'_{\psi_1}(-\mathbf{k}, -w)\}]$ corresponding to the noise terms $\eta_{\phi_1}$ and $\eta_{\psi_1}$ associated with the CC (Eq.(S11)) and TP equations (Eq.(S4)), respectively. The noise correlations are given by $< \eta'_{\phi_1}(\mathbf{k}, w)\eta'_{\phi_1}(-\mathbf{k}, -w) >= 2(k_a\phi_0 + k_b\phi_0^2 + D_\phi\phi_0 k^2)$, and $< \eta'_{\psi_1}(\mathbf{k}, w)\eta'_{\psi_1}(-\mathbf{k}, -w) >= 2D_\psi\psi_0 k^2$. The action $\mathcal{S}(\phi_1, \psi_1)$ in terms of $\phi_1(\mathbf{k}, w)$ and $\psi_1(\mathbf{k}, w)$ may be calculated using Eq.(S11) and Eq.(S4). The expression is too complicated to reproduce here, and is not needed for obtaining the main results. For the simpler case (for a single $\phi$ field) we have derived it elsewhere [2].

We obtain the following self-consistent equation for the self-energy $\Sigma_{\psi_1}(\mathbf{k}, \omega, \omega_\tau)$ from the calculation of response function:

$$\Delta\nu = \frac{D}{2\nu}\Sigma_{\psi_1}(\mathbf{k}, \omega, \omega_\tau) \tag{S22}$$

where, $\nu = D_\psi k^2 + \psi_0 k^2 U(\mathbf{k}) + \phi_0 k^2 U(\mathbf{k})$ and $D = 2D_\psi\psi_0 k^2$. Physically the self-energy term determines the contribution to the relaxation rate arising from the non-linearity. We can carry out the momentum count of Eq.(S22), keeping in mind that $\Delta\nu \sim k^z$, to extract the dynamic exponent $z$ in different time regimes.

#### II. SIMULATION MODELS:

In order to test the theoretical predictions and determine the mechanisms underlying the unusual dynamics, we simulated a three dimensional tumor spheroid with embedded TPs. An agent based model [9–11] is used for the tumor spheroid. The cells are treated as deformable objects. The size of the CCs increase with time as the tumor grows, and divide into two identical cells upon reaching a critical mitotic radius ($R_m$). The mean cell cycle time is $\tau$. The cell cycle time $\tau$ is expressed in units of $\tau = 15\ hrs$. The CCs can also undergo apoptosis. As in the theory, the TPs are inert, and their sizes and the number are constant throughout the simulations. We include CC-CC, CC-TP and TP-TP interactions. We use two potentials for the interactions.

##### A. Hertz potential

The form of the Hertz forces between the CCs is the same as in previous studies [9, 10, 12–14]. The physical properties of the CC, such as the radius, elastic modulus, membrane receptor and ligand concentration characterize the strength of the inter-cellular interactions. The elastic forces between two spheres with radii $R_i$ and $R_j$, is given by,

$$F_{ij}^{el} = \frac{h_{ij}^{\frac{3}{2}}}{\frac{3}{4}(\frac{1-\nu_i^2}{E_i} + \frac{1-\nu_j^2}{E_j})(\sqrt{\frac{1}{R_i} + \frac{1}{R_j}})}, \tag{S23}$$

where $E_i$ and $\nu_i$ are, respectively, the elastic modulus and Poisson ratio of the $i^{th}$ cell. Since, the CCs or the TPs are deformable, the elastic force depends on the overlap, $h_{ij}$, between two cells. The adhesive force, $F_{ij}^{ad}$, between the CCs is proportional to the area of contact ($A_{ij}$) [15], and is calculated using, [13],

$$F_{ij}^{ad} = A_{ij} f^{ad} \frac{1}{2}(c_i^{rec} c_j^{lig} + c_j^{rec} c_i^{lig}), \tag{S24}$$

where $c_i^{rec}(c_i^{lig})$ is the receptor (ligand) concentration on the surface of the cells, and are taken to be unity in the present study. The coupling constant $f^{ad}$ allows us to scale the adhesive force to account for variable receptor and ligand concentrations.

Repulsive and adhesive forces in Eqs.(S23) and (S24) act along the unit vector $\vec{n}_{ij}$ pointing from the centers of cells $j$ and $i$. Therefore, the net force on cell $i$ ($\vec{F}_i^H$) is given by the sum over its nearest neighbors [NN(i)],

$$\vec{F}_i^H = \Sigma_{j \epsilon NN(i)} (F_{ij}^{el} - F_{ij}^{ad}) \vec{n}_{ij}. \tag{S25}$$

To model the TP-TP and TP-CC interactions, we assume that the TPs are CC-like objects, which mimics experiments [16]. Therefore, CC-TP and TP-TP interactions are the same as CC-CC interactions.

##### B. Gaussian potential

In the theoretical treatment, we assumed that the CC-CC interaction is given by a sum of Gaussian terms (Eq. (S2)). For this potential, the force $\mathbf{F}_{ij}^G$ on cell $i$, exerted by cell $j$, is,

$$\mathbf{F}_{ij}^G = \frac{1}{(2\pi)^{3/2}} [\frac{\nu e^{\frac{-r^2}{2\lambda^2}}}{\lambda^5} - \frac{\kappa e^{\frac{-r^2}{2\sigma^2}}}{\sigma^5}] \mathbf{r} \tag{S26}$$

where $\mathbf{r}$ is $\mathbf{r}(i) - \mathbf{r}(j)$. We write $\lambda$ and $\sigma$ as $\lambda = \widetilde{\lambda}(R_i + R_j)$ and $\sigma = \widetilde{\sigma}(R_i + R_j)$, as the ranges of interactions corresponding to the repulsive and attractive interactions, respectively. In our simulations, the CCs grow and divide, their radii change in time, and therefore $\lambda$ and $\sigma$ also change in time. However, since these interactions are short-ranged, we assume them to be constant (as done in the theory). So, we fixed $\lambda = \widetilde{\lambda}(2R_d)$ and $\sigma = \widetilde{\sigma}(2R_d)$, where $R_d$ ($\approx 4\mu m$) is the size of a daughter cell (introduced in the next section). For simplicity, we write force $\mathbf{F}_{ij}^G = [\frac{\widetilde{\nu} e^{\frac{-r^2}{2\lambda^2}}}{\lambda^2} - \frac{\widetilde{\kappa} e^{\frac{-r^2}{2\sigma^2}}}{\sigma^2}]\mathbf{r}$, where $\widetilde{\nu} = \frac{1}{(2\pi)^{3/2}}\frac{\nu}{\lambda^3}$ and $\widetilde{\kappa} = \frac{1}{(2\pi)^{3/2}}\frac{\kappa}{\sigma^3}$. In the simulations, we fixed $\widetilde{\nu} = 0.03$, $\widetilde{\lambda} = 0.28$, $\widetilde{\kappa} = 0.003$ and $\widetilde{\sigma} = 0.4$.

#### C. Equation of Motion

The equation of motion governing the dynamics of TP and CCs is taken to be,

$$\dot{\vec{r}_i} = \frac{\vec{F}_i}{\gamma_i}, \tag{S27}$$

where $\dot{\vec{r}_i}$ is the velocity of $i^{th}$ CC or TP, $\vec{F}_i$ is the force on $i^{th}$ CC/TP (see equation S25 and S26), and $\gamma_i$ is the damping term (for details see reference [9]).

#### D. Cell division and Apoptosis

The CCs are either dormant or in the growth phase depending on the value of the pressure. The pressure on cell $i$ ($p_i$) due to $NN(i)$ neighboring cells is calculated using the Irving-Kirkwood equation,

$$p_i = \frac{1}{3V_i}\Sigma_{j\epsilon NN(i)}\mathbf{F}_{ij}\cdot d\mathbf{r}_{ij}, \tag{S28}$$

where $\mathbf{F}_{ij}$ is the force on the i$^{th}$ cell due to j$^{th}$ cell and $d\mathbf{r}_{ij} = \mathbf{r}_i - \mathbf{r}_j$. The volume of the $i^{th}$ cell ($V_i$) is $\frac{4}{3}\pi R_i^3$, where $R_i$ is the radius of the i$^{th}$ cell. If $p_i$ exceeds a pre-assigned critical limit $p_c$ ($= 1.7\times 10^{-6}$ MPa) the CC enters a dormant phase. The dormancy criterion serves as a source of mechanical feedback, which limits the growth of the tumor spheroid [17–21]. The volume of a growing cell increases at a constant rate, $r_V$. The cell radius is updated from a Gaussian distribution with the mean rate $\dot{R} = (4\pi R^2)^{-1}r_V$. Over the cell cycle time $\tau$,

$$r_V = \frac{2\pi(R_m)^3}{3\tau}, \tag{S29}$$

where $R_m$ is the mitotic radius. A cell divides once it grows to the fixed mitotic radius. To ensure volume conservation, upon cell division, we use $R_d = R_m 2^{-1/3}$ as the radius of the daughter cells. The resulting daughter cells are placed at a center-to-center distance $d = 2R_m(1-2^{-1/3})$ (Fig. 1 in the main text). The direction of the new cell location is chosen randomly from a uniform distribution on a unit sphere.

We initiated the simulations with 100 TPs and 100 CCs. The coordinates of the CCs and TPs were sampled using a normal distribution with mean zero and standard deviation

50 $\mu m$. The initial radii of the CCs and TPs were sampled from a normal distribution with mean 4.5 $\mu m$, and a dispersion of 0.5 $\mu m$.

#### III. DETAILS OF THE RESULTS

##### A. TPs exhibit sub-diffusive dynamics in the intermediate time regime

At $t < \frac{1}{k_b}$, the non-linear terms $\nabla \cdot \big(\psi_1(\mathbf{r},t)\int d\mathbf{r}'\psi_1(\mathbf{r}',t)\nabla U(\mathbf{r}-\mathbf{r}')\big)$ and $\nabla \cdot \big(\psi_1(\mathbf{r},t)\int d\mathbf{r}'\phi_1(\mathbf{r}',t)\nabla U(\mathbf{r}-\mathbf{r}')\big)$ describing the TP-TP and TP-CC interactions respectively, govern the scaling behavior of $\Delta_{TP}(t)$, the MSD. In the spirit of self-consistent mode coupling theory, we replace $\nu$ by $\Delta\nu$ in the self-energy term $\Sigma(k,\omega,\omega_\tau)$ (Eq.(S22)). Using the scale transformation, we find $\omega \sim k^z$, $\omega_\tau \sim k^{2z-2}$, $G_{\psi_1} \sim k^{-2z+2}$, $C_{\psi_1} \sim k^{-4z+4}$, and the vertex factor $V \sim k^z$. The relevant part $V$ is $\frac{1}{(D_{\psi_1}\psi_0 k^2)}(\{i\omega + D_{\psi_1}k^2 + \psi_0 k^2 U(\mathbf{k})\}\{(-\mathbf{k}'\cdot\mathbf{k})U(\mathbf{k}')\})$. The self energy term has the structure: $\Sigma(\mathbf{k},\omega,\omega_\tau) \sim \int \frac{d^d\mathbf{k}'}{(2\pi)^d}\frac{d\omega'}{2\pi}\frac{d\omega'_\tau}{2\pi}VVGC$. By carrying out the momentum count in $\Sigma(\mathbf{k},\omega,\omega_\tau)$, and noting that $\Delta\nu \sim k^z$, we find $\Sigma(\mathbf{k},\omega,\omega_\tau) \sim k^{d-z+4}$. Using Eq. (S22), we obtain $k^z \sim k^{d-z+4}$, which leads to $z = 2 + \frac{d}{2}$.

The MSD of the TPs scales with $t$ as,

$$< [r(t) - r(0)]^2 >\sim t^{2/z} = t^{\alpha_{TP}}. \tag{S30}$$

In the intermediate times, $\beta TP = \frac{2}{z} = 0.57$, a value that holds provided the interaction potentials involving the CCs and TPs are given by Eq. S2.

##### B. Hyper-diffusion of the TPs in the long time limit

At long times ($t \gg k_b^{-1}$), the effects of non-linearity in the TP-CC interactions together with non-equilibrium forces due cell division and apoptosis, determine the TP dynamics.

As in the previous section, we use the scale transformations: $\omega \sim k^z$, $\omega_\tau \sim k^{2z-2}$, $G_{\psi_1} \sim k^{-2z+2}$, $C_{\psi_1} \sim k^{-4z+4}$, $G_{\phi_1} \sim k^{-2z}$, $C_{\phi_1} \sim k^{-4z}$, and vertex factors $V_1 \sim k^2$, and $V_2 \sim V_3 \sim k^z$. The relevant part of $V_1$ is $\frac{1}{2(k_a\phi_0 + k_b\phi_0^2)}k_b(\mathbf{q}\cdot\mathbf{k})U(\mathbf{q})$, and the form of $V_2$ or $V_3$ is $\frac{1}{(D_\Psi \Psi_0 k^2)}\{i\omega + D_\Psi k^2 + \Psi_0 k^2 U(\mathbf{k})\}\{(-\mathbf{k}'\cdot\mathbf{k})U(\mathbf{k}')\}$. Noting that $\Delta\mu \sim k^z$, we find $\Sigma(\mathbf{k},\omega,\omega_\tau) \sim \int \frac{d^d\mathbf{k}'}{(2\pi)^d}\frac{d\omega'}{2\pi}\frac{d\omega'_\tau}{2\pi}V_1V_2V_3G_{\psi_1}G_{\psi_1}C_{\phi_1}C_{\psi_1} \sim k^{d+4-7z}$. From the self-consistent equation (Eq. (S22)) we find that the dynamic exponent, $z = (d+4)/8$. The value of $\alpha TP = \frac{2}{z} = \frac{16}{7} = 2.28$. Thus, we predict that the TPS must exhibit *hyper-diffusive* motion at long times.

##### C. k-dependent diffusion coefficients

One of our key findings is that the motion of the TPs is hyper-diffusive whereas the CC dynamics is super diffusive ($\alpha_{TP} > \alpha_{CC}$), whether the TPs are present or not. For

(a)

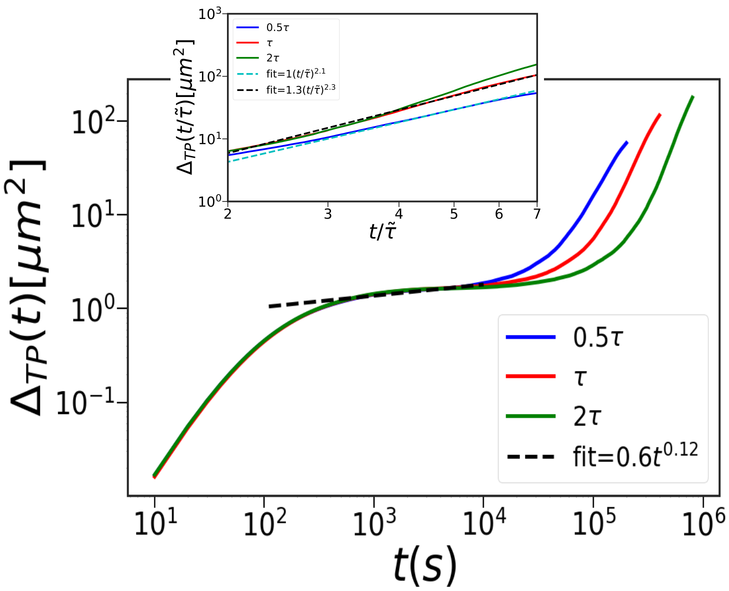


(b)

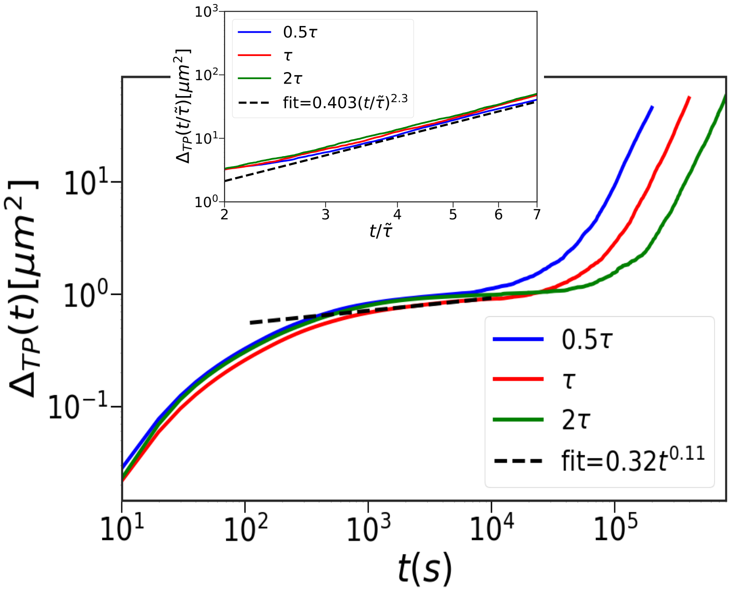


**FIG. S2:** MSD, $(\Delta_{TP}(t))$, of the TPs as a function of time $(t)$ for the two types of CC interactions. **(a)** $\Delta_{TP}$ calculated using Eq. $S25$ for the forces between the CCs and TPs. The curves are for 3 cell cycle times (blue – $0.5\tau$, red – $\tau$, and green – $2\tau$). Time taken to reach the super-diffusive regime, which is preceded by a sub-diffusive regime, increases as cell cycle time increases. In the long-time, $\Delta_{TP}(t)$ undergoes hyper-diffusive motion ($\Delta_{TP} \sim t^{\alpha_{TP}}$ with $\alpha_{TP} > 2$), which is highlighted in the inset. The x-axis of the inset plot is scaled by $\frac{1}{\tilde{\tau}}$. The black (cyan) dashed line shows exponent $\alpha_{TP} = 2.30$ (2.1). The curve with 0.5 $\tau$ is best fit best with $\alpha_{TP} = 2.1$. **(b)** Same as (a) except the Gaussian potential (equation S26) is used in the simulations. Interestingly, $\alpha_{TP}$ does not change appreciatively.

a homogeneous system, the density fluctuation obeys the Eq.(S4) without the non-linear terms. Thus, the equilibrium fluctuations, ($< \psi_1(k,t)\psi_1(k,0) >= \psi_0 \exp[-Dk^2t]$) decay exponentially. The diffusion-coefficient ($D$) is a constant, and the MSD exponent is unity. In this case, the relaxation time ($(Dk^2)^{-1} \approx k^{-z}$) with $z = 2$. Deviation from this standard situation is suggestive of anomalous diffusion. Systematic interactions and non-equilibrium forces between the CCs and TPs modify the density fluctuations, giving rise to an effective $k$-dependent diffusion coefficient.

Depending on the value of $z$, the TPs and CCs could exhibit sub or super or hyper diffusive motion. The non-linear term in Eq. (S4) for the TP-TP interactions renormalizes the diffusion coefficient $D$. The effective TP diffusion constant, $D_{TP} \sim k^{z-2} = k^{-9/8}$, and for CCs the diffusion coefficient $D_{CC} \sim k^{-5/8}$. The relaxation time for the dynamic structure factor for TPs ( $k^{-7/8}$) is small compared to the relaxation time for CCs ( $k^{-11/8}$), leading to a higher degree of anomaly in the diffusion of the TPs.

##### D. $\alpha_{TP}$ is nearly independent of the TP size

We varied the radius of the TP ($r_{TP}$) from $0.5r_c$ to $r_c$, where $r_c = 4.5\mu m$ is the average CC radius. Figure S4 shows $\Delta_{TP}(t)$ as function of $t$ for the Hertz potential (Eq. (S25)). Similar behavior is found for Gaussian potential as well. In the intermediate time regime, TPs with larger radius have higher MSD because they experience large repulsive forces due to higher excluded volume interactions. In the long time limit, $\Delta_{TP}$ exhibits hyper-diffusion (insets of Figure S4). The CC-TP interaction term,$\nabla \cdot \left(\psi_1(\mathbf{r},t) \int d\mathbf{r}'\phi_1(\mathbf{r}',t)\nabla U(\mathbf{r}-\mathbf{r}')\right)$, in Eq.(S4) shows that the radius only alters the interaction strength, and does not fundamentally alter the scaling behavior. The conclusion that the values of $\alpha_{TP}$ do not change, anticipated on theoretical grounds, is further supported by simulations.

##### E. Influence of the TPs on CC dynamics

We calculated $\Delta_{CC}(t)$ using the Hertz (Gaussian) potential as a function of the TP radius (Figure S5**a** (S5**b**). For both the potentials, the values of $\Delta_{CC}(t)$ for $t < \tau$ is greater as the TP sizes increase. Before cell division the number of CCs and TPs are similar, which explains the modest influence of the TPs on the dynamics of CCs in the intermediate time regime. The larger TPs undergo stronger repulsion (the repulsive interaction is proportional to $R^2$) initially, which increases the magnitude of $\Delta_{CC}$. The long-time dynamics is not significantly affected by the CC-TP interactions (see Figure S5). In the absence of the TPs, the CCs exhibit super-diffusion where the MSD scales as $t^{\alpha_{CC}}$ with $\alpha_{CC} = 1.33$. In the presence of the TPs, the CC dynamics remains super-diffusive with $\alpha_{CC} \approx 1.45$.

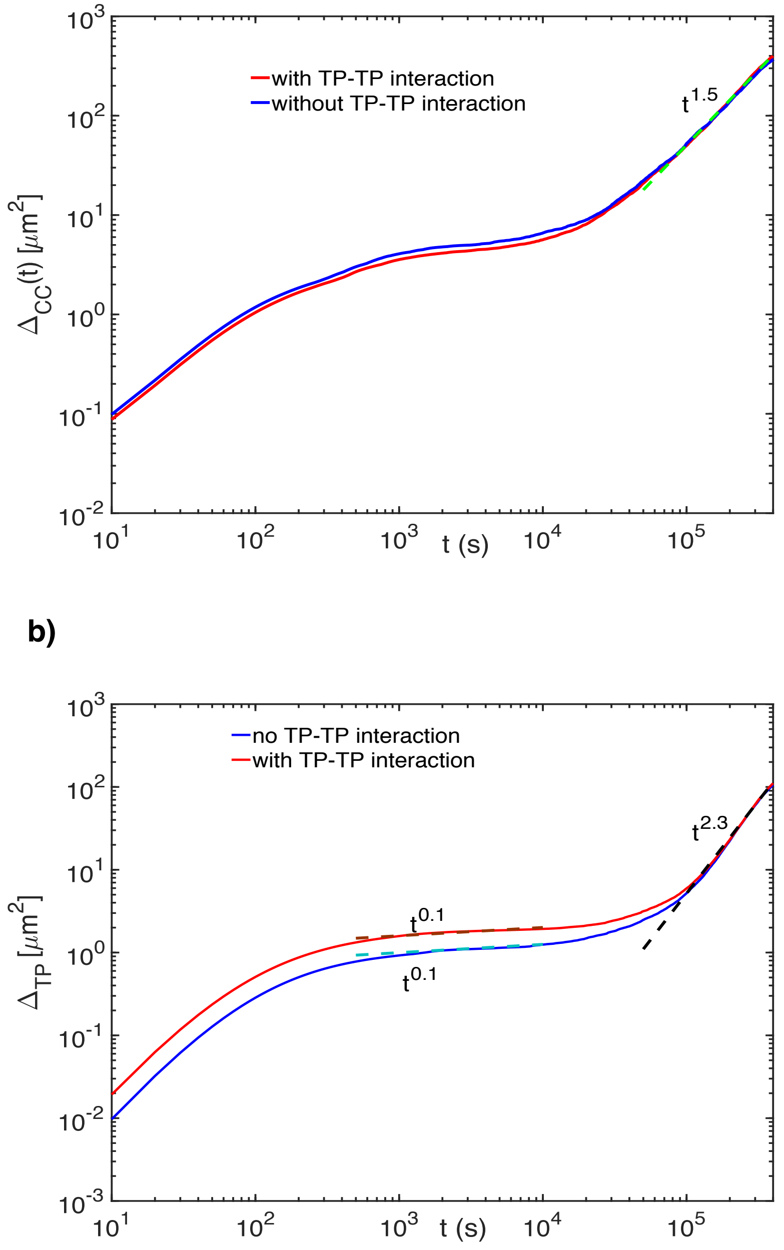


.

**FIG. S3: TP-TP interactions do not affect the long time dynamics of TPs or CCs.** (a) $\Delta_{CC}$ with (red curve) and without (blue) TP-TP interactions. The TP-TP interactions plays no role in the CC dynamics. The cyan dashed line shows $\alpha_{CC} = 1.5$ for both the cases. (b) $\Delta_{TP}$ with (red curve) and without (blue) TP-TP interaction. $\Delta_{TP}$ differs in magnitude in the intermediate time. However, TP-TP interactions plays no role in the long time dynamics of the TPs.

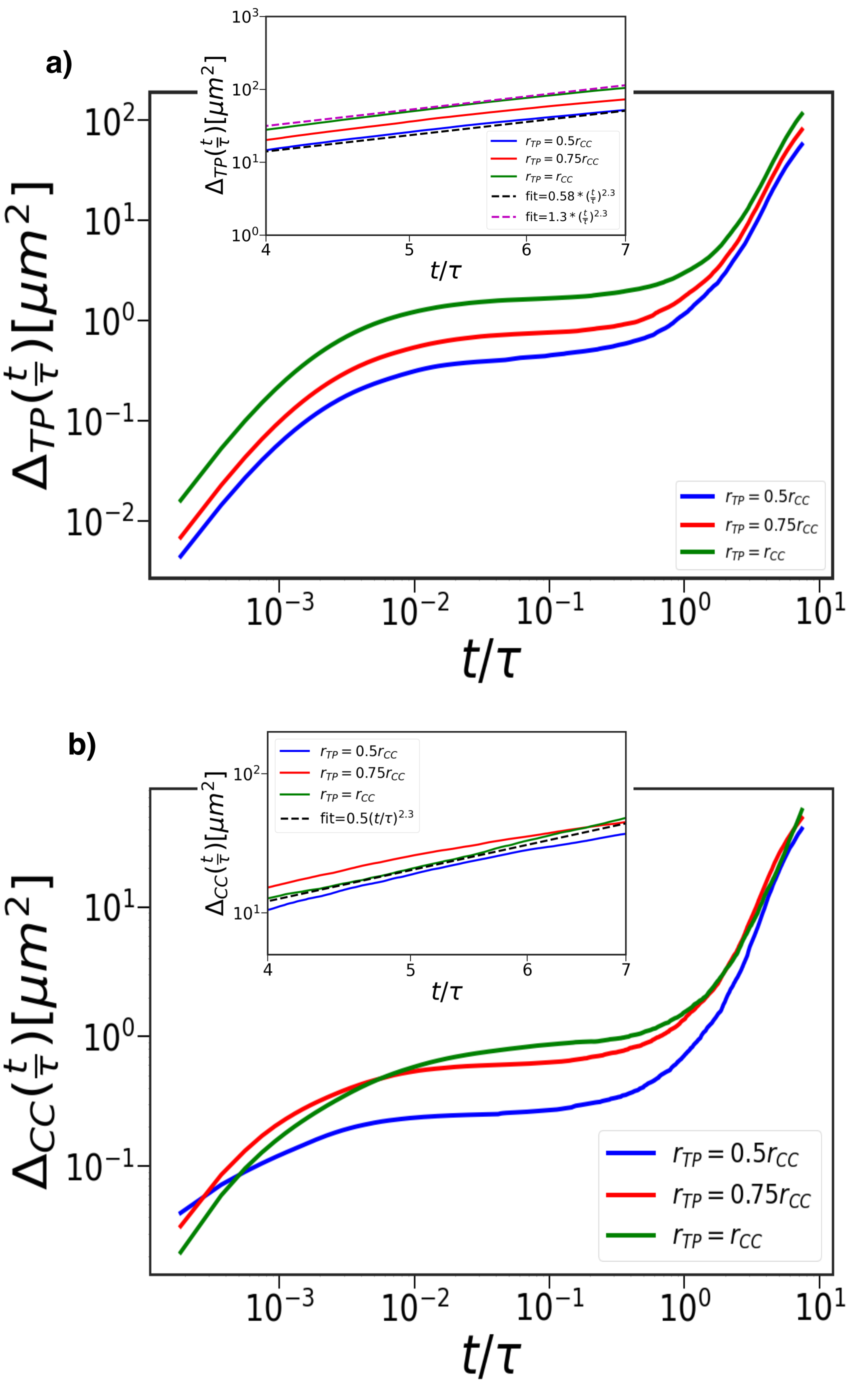


.

**FIG. S4:** Dependence of the TP size $\Delta_{TP}$ as a function of $t$. **(a)** Data are for the Hertz potential (see equation S23-S25). Time is scaled by $\tau$. From top to bottom, the curves correspond to decreasing TP radius ($r_{TP} = r_{CC}$ (green), $r_{TP} = 0.75r_{CC}$ (red) and $r_{TP} = 0.5r_{CC}$ (blue), where $r_{CC} = 4.5\mu m$ is average CC radius). TPs with larger radius have larger MSD values in the intermediate time ($\frac{t}{\tau} \leq \mathcal{O}(1)$). In the inset, we focus on the hyper-diffusive regime. The black and magenta dashed line serves as a guide to the eye with $\alpha_{TP} = 2.3$ **(b)** Same as (a) but with the Gaussian interactions.

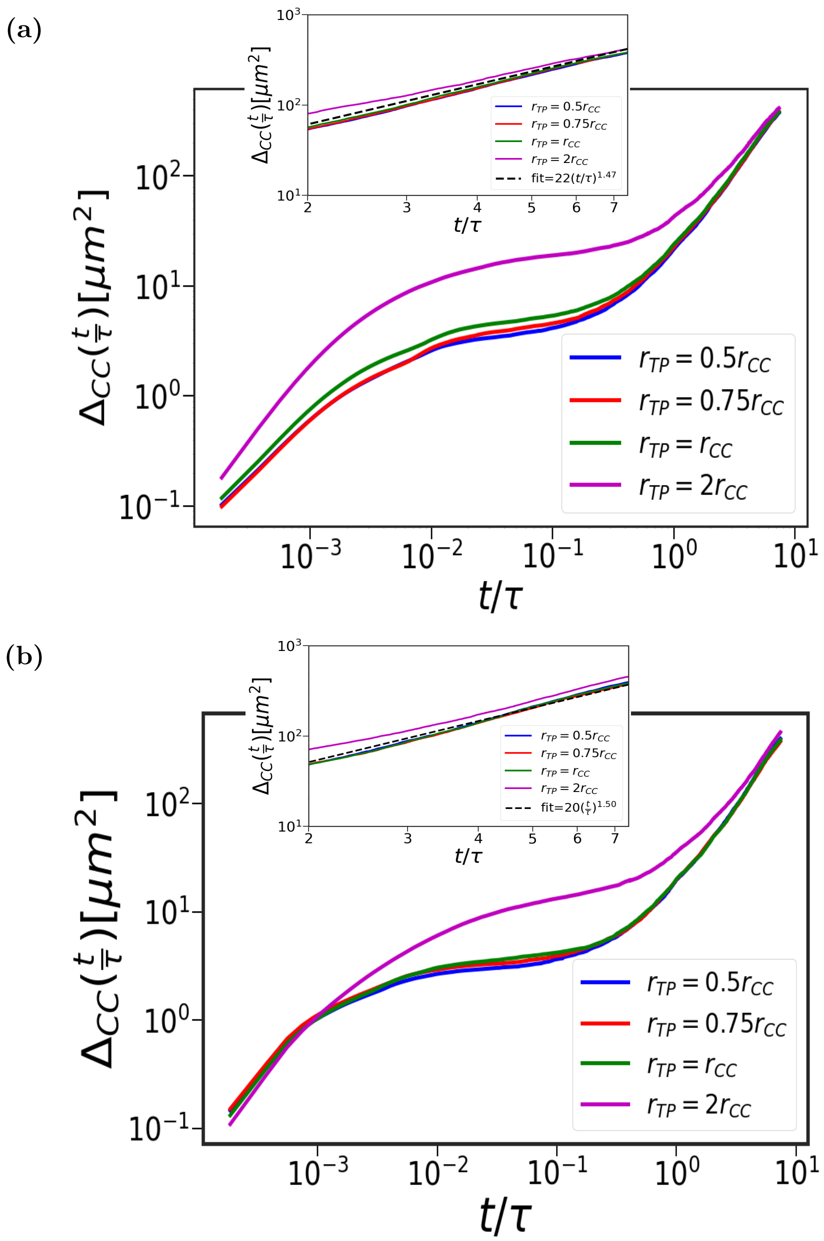


**FIG. S5:** Influence of TPs on CC dynamics using two potentials. $\Delta_{CC}$ as function of $t$ using the Hertz potential (equation S25). From top to bottom, the curves are for different values of the radius of the TPs (magenta $r_{TP} = 2r_{CC}$, green $r_{TP} = r_{CC}$, red $r_{TP} = 0.75r_{CC}$ and blue $r_{TP} = 0.5r_{CC}$ (appears to be hidden), where $r_{CC} = 4.5\ \mu m$ is the average cell radius). In the intermediate times, $\Delta_{CC}(t)$ is larger for TPs with larger radius. The inset focuses on the long time regime ($\frac{t}{\tau} > 1$). The black line is a guide to show the value of $\alpha_{CC} = 1.47$. (b) Same as (a) but with the Gaussian potential.

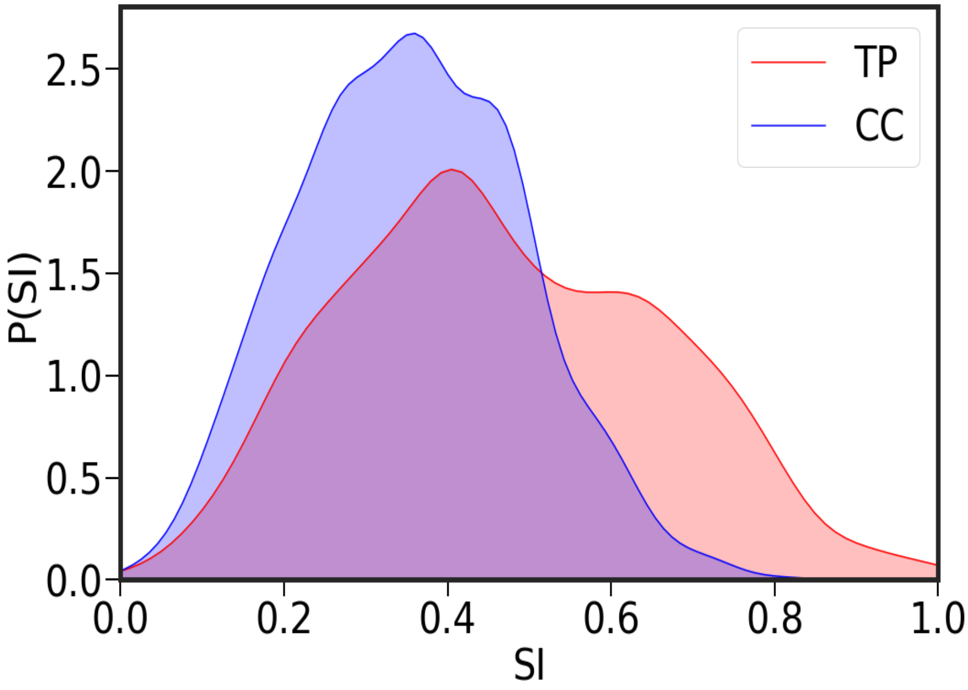


**FIG. S6:** Distribution of the Straightness Index (SI). The red (blue) plot for the TPs (CCs) shows that the TP trajectories are more rectilinear than the CCs.

##### F. Straigtness Index

The Straightness Index (SI) is, $SI_i = \frac{\mathbf{r}_i(t_f)-\mathbf{r}_i(0)}{\sum \delta \mathbf{r}_i(t)}$. The numerator $\mathbf{r}_i(t_f) - \mathbf{r}_i(0)$ is the net displacement of $i^{th}$ TP or CC, and the denominator $\sum \delta \mathbf{r}_i(t)$ is the total distance traversed. Fig. S6 shows that the TP trajectories are more straight or persistent over longer duration. During each cell division, the CCs are placed randomly causing the trajectories to be less persistent, thus explaining the decreased persistence in their motion.

[10] A. Malmi-Kakkada, X. Li, S. Sinha and D. Thirumalai, *arXiv preprint arXiv:1906.11292*, 2019.

[11] S. Sinha, A. Malmi-Kakkada, X. Li, H. Samanta and D. Thirumalai, *bioRxiv*, 2019, 842930.
